# Moisture and competition constrain ephemeral resource quality for burying beetle reproduction

**DOI:** 10.1101/2024.11.04.621899

**Authors:** Tracie E. Hayes, Léo Lassérès, Louie H. Yang

**Affiliations:** Department of Entomology and Nematology, and the Center for Population Biology, University of California, Davis CA 95616 USA; École d’ingénieurs de Purpan, 75 Voie du Toec, 31076 Toulouse, France

**Keywords:** ephemeral resources, climate change, scavengers, moisture, competition, burying beetles, fog decline

## Abstract

Shifts in abiotic factors such as temperature and moisture can change the availability of resources, especially under climate change. Both abiotic and biotic drivers can have profound, rapid effects on species distribution, survival, and reproduction. Little is known about how abiotic factors affect the availability of ephemeral resources. Burying beetles (*Nicrophorus* spp.) are specialist users of ephemeral resources, as their reproduction requires locating, defending, and burying a small carcass. Environmental moisture, such as coastal fog, could change how quickly carcasses dry out. We tested the role of carcass moisture and interspecific competition with a generalist scavenger, *Heterosilpha* spp., on reproduction by placing pairs of *Nicrophorus guttula* in field chambers with fresh and experimentally dehydrated mouse carcasses. Pairs that were given fresh mouse carcasses were more likely to carry out reproductive behaviors and produce viable offspring than pairs that were given a partially dehydrated mouse. For those pairs that reproduced, competition limited the number of offspring. These results indicate that shifts in abiotic factors under climate change, along with biotic factors like competition, can reduce the availability of ephemeral resource patches for consumers.

## Introduction

Considering the effects of both biotic and abiotic drivers on the success of species has been a goal of ecologists as they attempt to predict biodiversity shifts under global change (Morris et al. 2020). For example, more variance in the temporal occupancy of North American bird species is explained by environmental variables compared to competitor abundance, but for migrants with small ranges, competitor abundance is a stronger driver (Snell Taylor et al. 2020). Similarly, bumble bees are sensitive to both biotic and abiotic drivers depending on life history stage (Ogilvie and CaraDonna 2022), and across 208 terrestrial plant species, biotic and abiotic drivers have effects of a similar strength on population growth rate (Morris et al. 2020). Though species distribution models have primarily focused on abiotic drivers, both abiotic and biotic must be considered in a global change context (Morris et al. 2020).

The shifting of temperature and moisture in many environments is one of the direct effects of climate change (IPCC 2023). Abiotic drivers are changing rapidly, and the effects of these changes are still unknown in many systems. This provides an opportunity to test the effects of certain abiotic drivers that we know are changing in particular places, to project how continued change might affect a species. These manipulations, alongside relevant manipulations of biotic drivers, can set up experiments that help shed light on the relative and interactive effects of these two drivers under a changing climate.

Fog is a potentially important abiotic driver in coastal ecosystems that would otherwise be dry during the summer. However, reduced summertime fog is a likely outcome of climate change in many ecosystems. For example, coastal prairies are becoming drier in the summers as fog decreases along the northern California coast (Johnstone and Dawson 2010, Torregrosa et al. 2014). This could have implications for the success of species in these habitats that rely on the moisture provided by summertime fog (Karban and Huntzinger 2018, 2021).

Ephemeral resource patches may be especially vulnerable to changes in abiotic factors and can provide testable systems to study the effects of abiotic and biotic drivers. Ephemeral resource patches are a combination of Elton’s “minor habitats” (Elton 1949) and resource pulses (Yang et al. 2008, 2010), as they are repeated, smaller magnitude resource pulses within a spatial context (Butterworth et al. 2022). Species that rely on ephemeral resource patches can be useful systems when testing for the effects of abiotic and biotic drivers, as it is often easy to manipulate resources when they are small and only available to consumers for a short period of time. The resource can be manipulated to mimic levels of the key abiotic factor, and replicates can be acquired over a relatively small area with realistic field conditions. These species may be especially sensitive to changes in their resources due to both abiotic and biotic shifts, such as desiccation and competition, respectively (Butterworth et al. 2022). Further compression of the window of time in which an ephemeral resource is available could have harmful effects on consumers, making these consumers indicators of change that might otherwise be missed.

We tested the effects of resource moisture and competition on reproductive success of the burying beetle *Nicrophorus guttula* in the field. Burying beetles are specialists on ephemeral resource patches as they require a small carcass that they defend, bury, and protect in order to raise their young belowground (Scott 1998). Authors of recent studies have measured and tested how multiple species of *Nicrophorus* coexist at the same sites. Depending on the location, they may or may not partition resources in space via habitat preferences (Garfinkel and McCain 2023, Burke et al. 2024), or time via differences in seasonal activity (Wettlaufer et al. 2021). In this study, we ask how increased dessication affects the reproductive success of a single species.

These types of shifts in resource quality could be taking place across a wide range of habitats and seasons under the new normal of constant global change, so they have relevance across the resource partitioning studies just mentioned.

We hypothesized that carcasses are drying out faster as summertime fog decreases along the northern California coast (Johnstone and Dawson 2010, Torregrosa et al. 2014), and that this increased desiccation may constrain the reproductive success of burying beetles. We compare this abiotic effect with the biotic effect of competition, with conspecifics, congeners, other carrion beetles, other invertebrate scavengers (e.g. blowflies), and vertebrate scavengers (Wilson et al. 1984, Trumbo 1990, Scott 1998, Trumbo and Bloch 2002, Eggert et al. 2008). We also considered the effects of phoretic mites that may act as resource competitors or even predators of beetle eggs (Wilson and Knollenberg 1987, Scott 1998), but can also act as mutualists by consuming the eggs of blowflies, a common resource competitor (Sun and Kilner 2020). These biotic interactions are summarized in Figure 1A.

**Figure 1.**
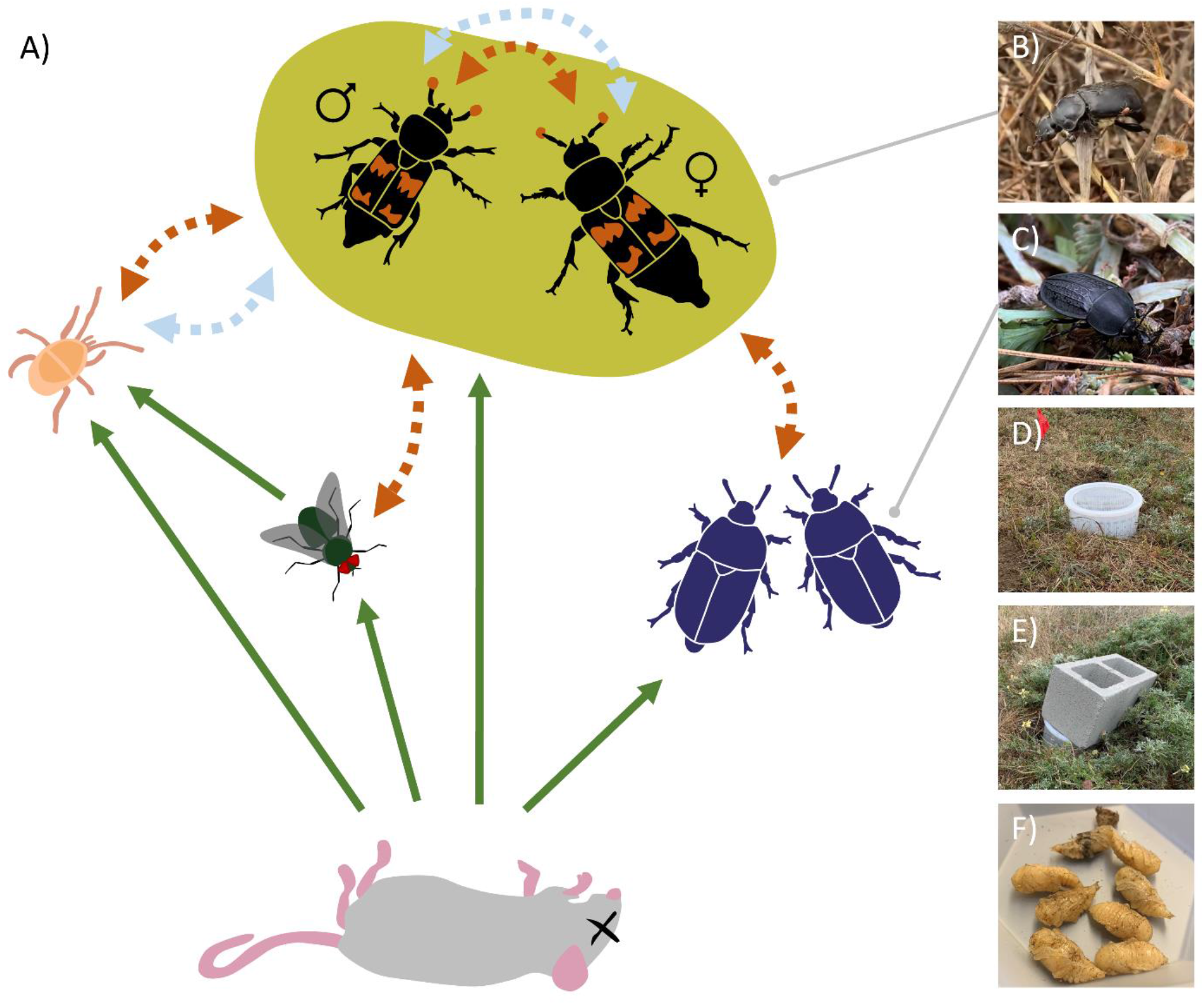
A) A possible network of burying beetles (top), a carcass resource (bottom), and multiple interacting species (left, a phoretic mite and blowfly; right, a confamilial competitor, *Heterosilpha* spp.). Straight green arrows represent consumption; double-sided, dark orange dotted errors represent potential competition or negative interactions; and double-sided light blue dotted arrows represent potential mutualism or facilitative interactions. Note that species are not to scale. See text for more information. B) *Nicrophorus guttula* with a phoretic mite. C) *Heterosilpha* spp.. D) Experimental chamber, halfway submerged in the soil, with a lid made of a layer of both wire mesh and fine mesh. E) A cinder block is placed over experimental chambers. F) Collected pupae from an experimental chamber that had successful reproduction.

In this experiment, we tested the effects of resource moisture and competition from a generalist carrion beetle (*Heterosilpha* spp.) on the reproduction of the burying beetle *N. guttula*, a different species that specializes on ephemeral resources. We asked: How does reduced resource moisture and increased competition affect the reproduction of *N. guttula*? We predicted that pairs that received fresh carcasses would have more successful reproduction than those that received dried carcasses. We also predicted that pairs that did not experience competition would show greater reproduction. Finally, we predicted that the combination of both reduced resource moisture and increased competition would limit reproduction the greatest.

## Material and methods

### Site

We conducted this experiment at the Bodega Marine Reserve in Bodega Bay, California (38.319061, -123.072443) between June and September 2022. The site is composed of coastal prairie, dunes, and rocky intertidal habitat. Summers are mild (typical daily means 10-15 C, with occasional periods of 16-18 C), with fog often present early in the mornings (mean relative humidity 90-100%) (WRCC 2024). The coastal prairie is primarily composed of species such as bush lupine (*Lupinus arboreus*), seaside daisy (*Erigeron glaucus*), woolly sunflower (*Eriophyllum* spp.), and ice plant (*Carpobrotus* spp.) (Barbour et al. 1973).

### Trapping beetles

We set up pitfall traps baited with mouse carcasses in the coastal prairie to collect *N. guttula* and *Heterosilpha* spp. for experiments (Figure 1B, 1C). Most of the *Heterosilpha* beetles were likely to be *H. ramosa*, as *H. aenescens* is rare at this site. We caught most of the beetles used in this experiment along a transect of 13 traps installed 12.5 meters apart on the coast bluffs south of the Bodega Marine Lab. When collecting *N. guttula*, we sexed them, counted the number of phoretic mites currently attached, and took photographs to measure size in ImageJ. We usually visited traps in the mornings to collect the beetles that would then be placed into experimental chambers later that day.

### Mouse carcass preparation

We purchased large frozen mice (mean = 21 g, range = 17-26 g, SD = 2 g) from a commercial source (Gourmet Rodent via Petco.com
). We dried half of the mice in a food dehydrator for 5-6 hours at 45 C. On average, dried mice lost 25% (SD = 4%) of their weight in water. This corresponds to between one and one and a half July days of mouse carcasses naturally drying out and decomposing at the site (Figure S1). Mice carcasses also tend to lose more water weight (dry out faster) during less humid and less foggy periods at the Bodega Marine Reserve (Figure S2). All mice (dried and undried) were refrozen to minimize further changes until they were placed into experimental chambers, where they thawed rapidly.

### Setting up experimental chambers

We set up 63 experimental chambers (Figure 1D) between June 22 and August 10, 2022. We used 5-liter buckets (16 cm tall by 22 cm diameter) with modified lids that had a layer of wire mesh with quarter-inch openings and a layer of fine mesh, to prevent beetles from getting in or out. A cinder block was placed over each chamber to prevent vertebrate disturbances (Figure 1E). Each bucket was filled with local soil and plants and buried to a depth of 8 cm. We placed experimental chambers along a series of transects in the coastal prairie, approximately 25 meters apart. At each transect point, we haphazardly placed experimental chambers by throwing a small object into the air and digging the hole wherever the object landed.

Each experimental chamber received a male-female *N. guttula* pair. We randomly assigned chambers to receive one of four treatments: (1) 14 received an undried carcass and no *Heterosilpha* beetles, (2) 16 received an undried carcass and a male-female pair of *Heterosilpha*, (3) 16 received a partially dried carcass and no *Heterosilpha* beetles, and (4) 17 received a partially dried carcass with a male-female pair of *Heterosilpha*.

### Measuring reproductive output

We checked each chamber after about 39 days (range = 35-42 days, SD = 2), between August 6, 2022, and September 19, 2022, when the adult beetles that we started the experiment with had sufficient time to reproduce. We counted all larvae and pupae (Figure 1F). We also noted if the carcass was buried, even if there were no larvae or pupae present.

### Analysis

We tested two models to analyze these data. First, we analyzed the role of carcass dryness and competition from *Heterosilpha* on binomial reproductive success with a binomial generalized linear model (GLM) using the R packages lme4 and jtools (Long 2023, Bates et al. 2023). We also included the initial undried mass of the carcass, day of year, female size, male size, and total number of mites on adults as covariate predictors. The initial undried mass of the carcass was included to control for initial variation in carcass size. Day of year that the experimental chamber was set up was included to control for seasonality. Female and male size were included to account for effects of size on fecundity and/or competition between mates during the reproductive process. Total number of mites on adults was included to account for effects of the mites as either competitors or mutualists.

Second, we made another GLM with a Poisson distribution with a smaller dataset that only included experimental chambers that had successful reproduction. The model had number of offspring as the response variable, with the same predictors as the previously mentioned model.

## Results

Partial (approximately 25%) carcass desiccation decreased the likelihood of reproduction by nearly 18-fold (χ^2^ = 27.29, df = 1, p < 0.001, Figure 2A and 2B). While 53% of pairs with fresh carcasses reproduced, only 3% of pairs (1 of total) with partially dried carcasses reproduced. Successfully reproducing pairs had on average carcasses with a fresh weight 0.98 g heavier than those pairs who did not reproduce (χ^2^ = 5.42, df = 1, p = 0.02, Figure 2B). The number of offspring per pair was not affected by carcass dryness (χ^2^ = 0.66, df = 1, p = 0.42) or carcass fresh weight (χ^2^ = 0.0005, df = 1, p = 0.98, Figure 2B). We found that all pairs given a fresh mouse buried it, even if they did not reproduce (Figure 3). Pairs given a dry mouse only buried it 29% of the time (Figure 3).

**Figure 2.**
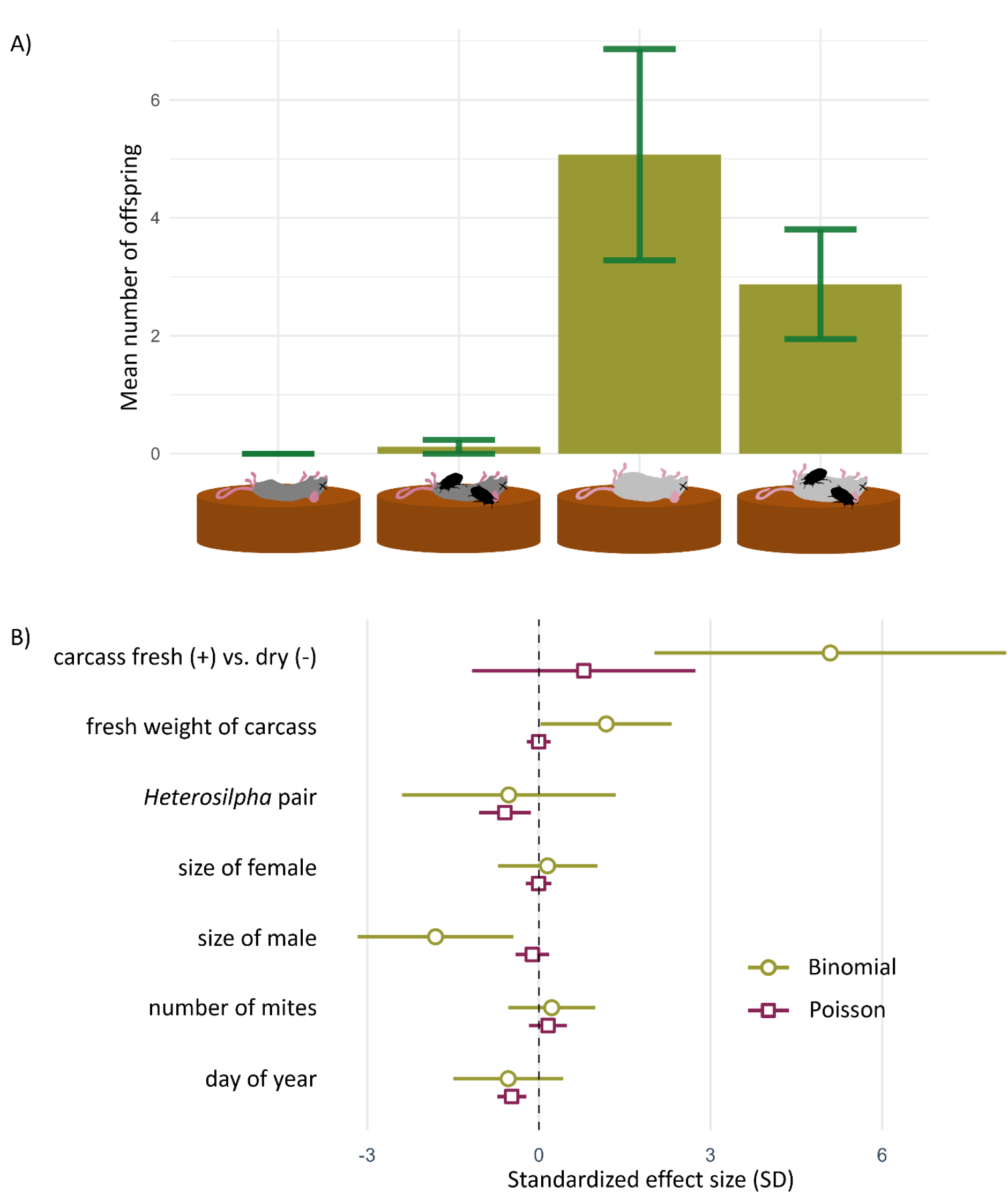
A) Mean number of offspring with standard error bars, for each treatment (dry mouse, dry mouse + competition, fresh mouse, fresh mouse + competition). B) Scaled effect size plot with standard error for each variable in the GLM with a binomial distribution (likelihood of reproduction, green) or in the GLM with a Poisson distribution (number of offspring, purple).

**Figure 3.**
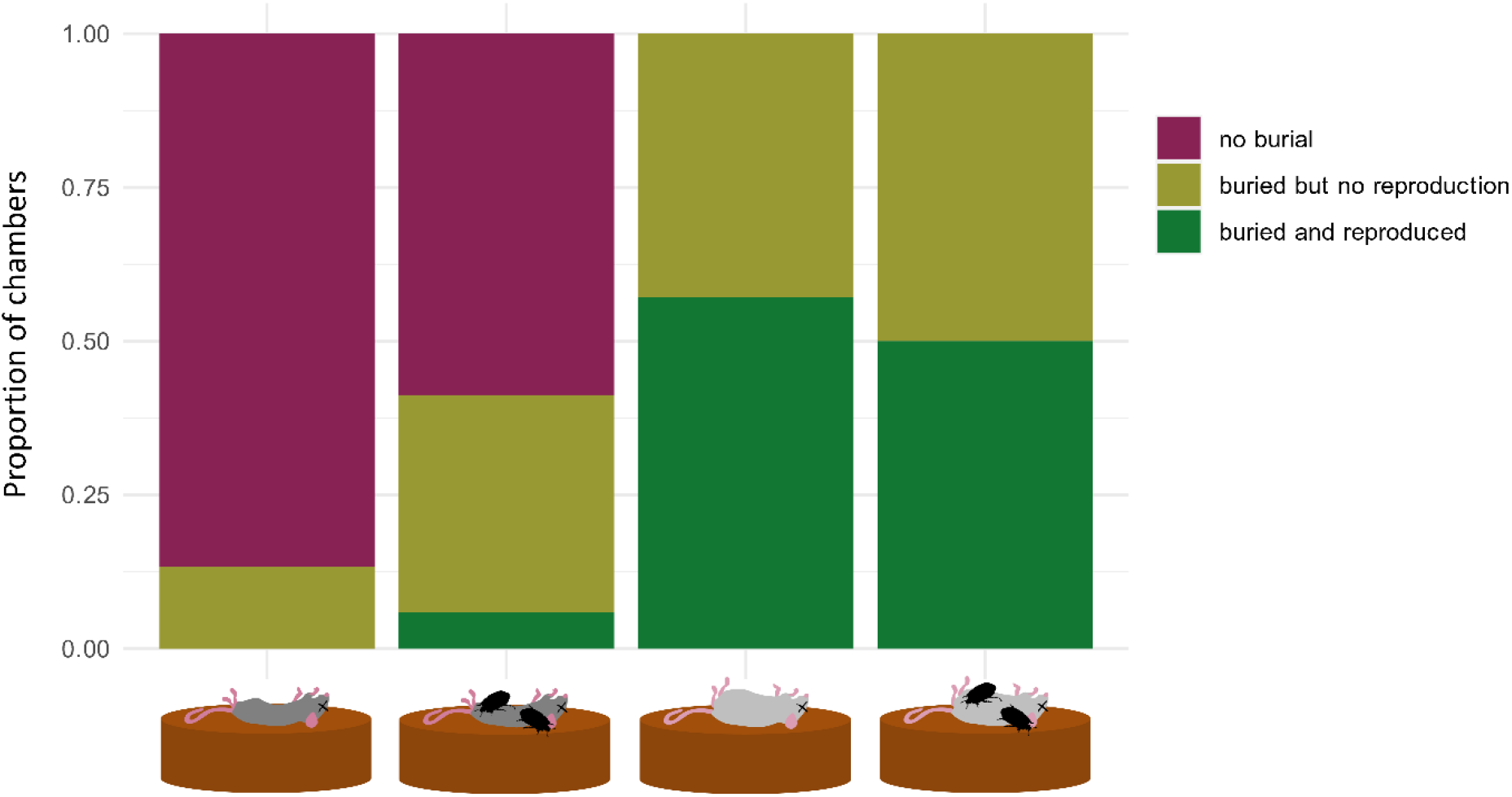
Proportion of experimental chambers within each treatment group (dry mouse, dry mouse + competition, fresh mouse, fresh mouse + competition) that had carcasses that were not buried (purple), buried with no reproduction (light green), or buried with reproduction (dark green).

Competition from *Heterosilpha* had no effect on the likelihood of reproduction, with pairs with and without competition reproducing about 27% of the time (χ^2^ = 0.31, df = 1, p = 0.58, Figure 2B). For pairs that did reproduce, however, receiving a pair of *Heterosilpha* competitors resulted in an average of 3.5 fewer offspring compared to *N. guttula* pairs that did not receive an *Heterosilpha* pair (χ^2^ = 7.08, df = 1, p = 0.01, Figure 2B). Of the pairs that reproduced, the average number of offspring was 7.0 (range = 1-18, SD = 5.3). The average brood size was 5.1 (SD = 6.7) for all pairs that received a fresh mouse and no competition, and 2.9 (SD = 3.7) for all pairs that received a fresh mouse with competition (Figure 2A).

Smaller males increased the likelihood of reproduction (χ^2^ = 12.61, df = 1, p < 0.001), but female size had no effect (χ^2^ = 0.12, df = 1, p = 0.73, Figure 2B). For each millimeter increase in male pronotum width, there was a 90% decrease in the likelihood of reproduction. Male and female size had no effect on number of offspring (male: χ^2^ = 0.57, df = 1, p = 0.45; female: χ^2^ = 0.003, df = 1, p = 0.96; Figure 2B). Though it appears that more phoretic mites might have increased the number of offspring marginally, number of mites had no effect on the likelihood of reproduction (χ^2^ = 0.36, df = 1, p = 0.55) or the number of offspring per pair (χ^2^ = 0.90, df = 1, p = 0.34, Figure 2B). Pairs were more likely to have more offspring earlier in the season, as each additional day added to the date of setting up the chamber resulted in a 4% decrease in offspring count (χ^2^ = 13.93, df = 1, p < 0.001). Day of year the experimental chamber was set up had no effect on likelihood of reproducing successfully (χ^2^ = 1.25, df = 1, p = 0.26, Figure 2B).

## Discussion

Our finding that carcass dryness strongly inhibits burying beetle reproduction (Figure 2A, 2B) may have implications for the population ecology of these beetles in increasingly arid regions. Our results suggest that the window for successful reproduction on a carcass could be substantially shorter under increasingly dry conditions, reducing opportunities for burying beetle reproduction.

Abiotic changes may have especially strong effects on the consumers of ephemeral resource patches. Since burying beetles are specialists of small carcasses, there is already a short window of opportunity during which the resource must be found and monopolized to reproduce successfully. Our findings are consistent with the recent decline of meadow spittlebugs, a species that is dependent on moist refugia along the coast, during a period with increasingly dry summers (Karban and Huntzinger 2018, 2021). In our study, the finding that 97% of burying beetle pairs were unable to reproduce with a 25% dehydrated carcass indicates a strong aversion to desiccated carcasses.

Carcass dryness inhibited both burial and reproduction outright, while competition appeared to limit the number of offspring for reproducing pairs. This suggests that environmental moisture could affect the availability of usable carcasses over the landscape, while competition reduces reproductive output at a smaller spatial scale once a usable carcass has been secured.

We consider a few possible reasons for the relatively small effect of competition in our study (Figure 2B). Previous studies have not looked at the long-term effects of competition on reproductive output, especially after a time span of 39 days in which burying, fighting, parental care, larval movement, and pupation have potentially occurred (Wilson et al. 1984, Trumbo 1990, Scott 1998, Trumbo and Bloch 2002, Eggert et al. 2008). Burying beetles have multiple adaptations making them very effective competitors, including their burying behavior to conceal carcasses, cooperation between mates, and their fighting abilities (Scott 1998). Because of this, we might not expect large effects of the presence of *Heterosilpha* on reproductive success. However, *Heterosilpha* is attracted to carrion for feeding and reproducing near it, so even if the *N. guttula* pair is not expending energy fighting off the competitors, they may lose some of the carcass mass to *Heterosilpha* feeding.

We found that the effects of moisture outweighed the effects of variables that have been previously shown to affect reproductive output, such as carcass size (Wilson and Fudge 1984, Scott 1998, Hopwood et al. 2016) (Figure 2B). Though we did not include a large range in carcass size (range = 17-26 g, SD = 2), we believe this range is realistic to what is available to these burying beetles.

Smaller males were more likely to reproduce (Figure 2B). This result could underestimate the effect of male size because this experiment did not require the pair to protect the carcass from congeners and conspecifics (Otronen 1988, Bartlett and Ashworth 1988, Safryn and Scott 2000). However, this result could also reflect the fact that smaller males might require less food during the preparation of the carcass, leaving more for the female and offspring (Hopwood et al. 2016). If the female deemed the male no longer useful in the reproductive process, smaller males also might be easier for the female to kill than larger ones.

There were no significant effects of the number of phoretic mites, but the nonsignificant effects were positive (Figure 2B). When blowflies are particularly productive, phoretic mites can benefit their host by removing blowfly eggs from the carcass (Sun and Kilner 2020). Phoretic mites can also help in competitive interactions between burying beetle individuals if there are enough to sufficiently warm up the beetle for battle (Sun et al. 2019). In our experiment, it is possible that having a greater number of mites aided in cleaning the carcass of unwanted competitors that may have been able to get in through the mesh. Even though adult flies are unable to get through the mesh, they are able to deposit eggs across the air gap (Yang 2006). We noted dipteran pupal cases (empty or full) four times when checking for reproduction.

We show that moisture can have a profound effect on how an ephemeral resource is received by its consumer. These results suggest that desiccation could limit the resources available for burying beetle reproduction. This increased scarcity may also increase the effects of competition and its additional negative effects on reproduction. The ability of drier and warmer conditions to exacerbate resource scarcity in space and time is a largely unexplored area in ecology. Future research concerning the effects of climate change on resource scarcity, and the resulting effects on species across the specialist-to-generalist continuum, would be helpful in determining which species will be especially sensitive under continued global change.

