## Supplemental Figures for "Moisture and competition constrain ephemeral resource quality for burying beetle reproduction"

Tracie E. Hayes[1*]

Léo Lassérès[2]

Louie H. Yang[1]

[1]Department of Entomology and Nematology, and the Center for Population Biology, University of California, Davis CA 95616 USA

[2]École d'ingénieurs de Purpan, 75 Voie du Toec, 31076 Toulouse, France

**Supporting Information**

Figure captions:

Figure S1. Remaining carcass weight over time, with each line representing a single mouse carcass placed out in the field at the Bodega Marine Reserve. Mice were secured under a small cage of mesh and hardware cloth and able to experience the elements. They were weighed in the field twice a day on July 2-6, or July 18-22, 2024. Each time period started with a unique set of mouse carcasses weighing between 17-25g that had been thawed over the previous 24 hours. Lines cut off before the end of the time period were either buried by burying beetles who got in or stolen by vertebrate scavengers.

Figure S2. Each point represents an interval of time over which a mouse carcass experienced a weight change. The y-axis indicates the rate of carcass weight change, so the number of grams lost (or gained) over the interval divided by the length of the interval in hours. The x-axis in the top panel indicates the mean relative humidity over that interval, calculated by summing the mean relative humidities for each hour within the interval divided by the length of the interval in hours. The x-axis on the bottom panel indicates the rate of "bucket tips" (pulses, or a measure of fog accumulation) per hour within the interval divided by the length of the interval in hours.

Figures:

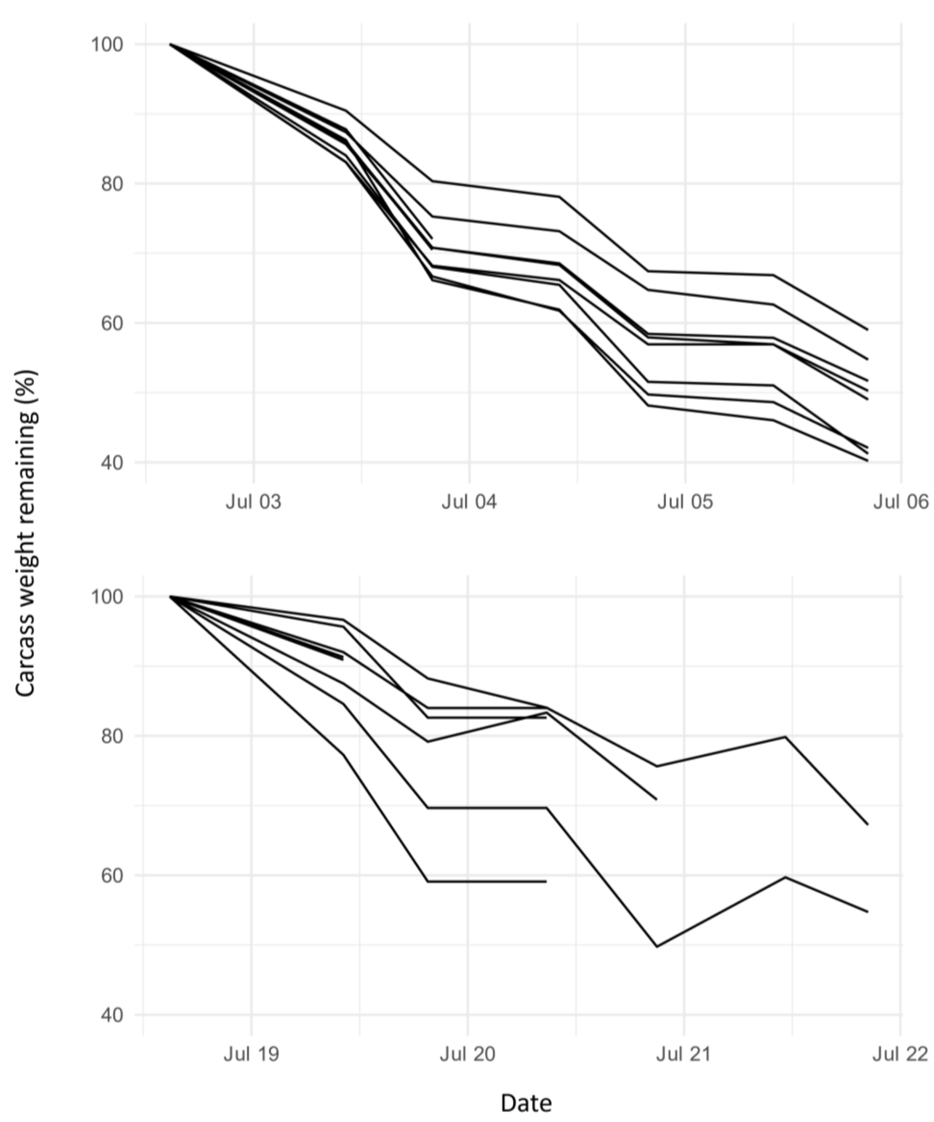


Figure S1

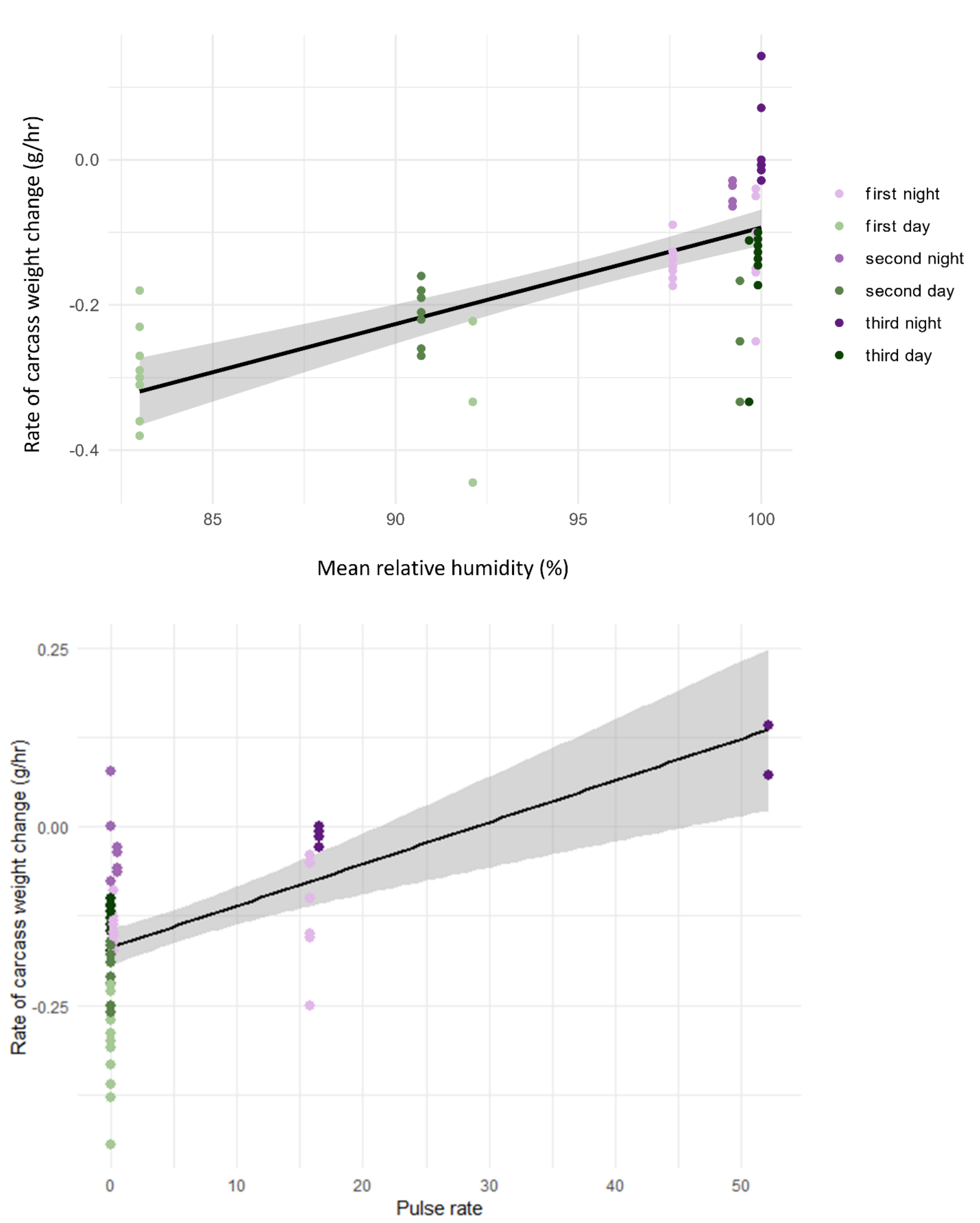
Rate of carcass weight change (g/hr)
0.0
-0.2
-0.4
85
90
95
100
Mean relative humidity (%)
first night
first day
second night
second day
third night
third day
Rate of carcass weight change (g/hr)
0.25
0.00
-0.25
0
10
20
30
40
50
Pulse rate

Figure S2
